# Elucidating Protein Dynamics through the Optimal Annealing of Variational Autoencoders

**DOI:** 10.1101/2025.01.08.632051

**Authors:** Subinoy Adhikari, Jagannath Mondal

## Abstract

Proteins traverse intricate conformational landscapes, with transitions and long-lived states that hold the key to their biological function. Yet, unraveling these dynamics remains a formidable challenge. An emerging approach has been to train the conformational ensemble via deep Variational autoencoders (VAEs) in a bid to machine learn the underlying reduced dimensional representation. However, training VAEs typically involves a fixed *β* value of 1, where *β* acts as the crucial weighing factor between the reconstruction and regularization terms. This static setup can often lead to posterior collapse, which significantly hinders the model’s ability to capture complex protein dynamics accurately. To mitigate this issue, annealing the *β* parameter offers a potential alternative. However, this approach frequently falls short in fully addressing the problem, majorly due to arbitrary choice of upper bound of *β* and annealing schedule. In this work, we introduce an innovative approach for selecting the *β* parameter by utilizing the Fraction of Variance Explained (FVE) score to identify its optimal value. We demonstrate that training annealed VAEs at their optimum *β* in a single cycle consistently outperformed their non-annealed counterparts, as evident from their higher Variational Approach for Markov Processes-2 and Generalized Matrix Rayleigh Quotient scores and distinct free energy surface minima on both folded and intrinsically disordered proteins. The improved latent space representations significantly improve state space discretization, thereby refining Markov State Models and providing more accurate insights into conformational landscapes as reflected in distinct contact maps. These findings not only underscore the potential of annealed VAEs in resolving complex conformational spaces but also highlight the critical interplay between annealing schedules and latent space structures.

---

Dimension reduction is a crucial method in understanding long molecular dynamics (MD) data. For a protein with N atoms, the system’s dimensionality is 3N. This high dimensionality makes it challenging to identify conformational changes such as domain closure or flexible loop movements^1^ which may be lost in the small rapid fluctuations of each atom in the high dimension space. High dimensionality includes both short-term fluctuations and long-term conformational changes, which can mask the identification of stable, long-lived states. ^2,3^ In the full 3N-dimensional space, tracking and interpreting binding^4^ and unbinding events,^5^ allosteric effect,^6^ or induced fit mechanisms in response to a ligand^7^ or protein partner can be obscured by the vast number of atomic positions. These interactions often cause specific, localized changes that may go unnoticed in high-dimensional space. Therefore, there is a need to better represent the data in a fewer number of dimensions. Several dimension reduction methods have been actively employed to preserve the key properties and changes in the original space.^8^ For instance, Principal Component Analysis (PCA)^9^ and time-lagged Independent Component Analysis (tICA)^10 11^ are popular for capturing variance and slow motions but are limited by their linearity. Nonlinear methods like t-SNE (t-distributed Stochastic Neighbor Embedding)^12 13^ and UMAP (Uniform Manifold Approximation and Projection)^14^ excel at revealing complex, nonlinear patterns in protein dynamics.

More recently, deep learning based approaches such as autoencoders^15–17^ have gained popularity over other non linear dimension reduction methods. An autoencoder compresses the input data to a latent space, which typically contains a few dimensions, followed by its decompression to reconstruct the input data. The training objective is to minimize a loss function, ensuring that the latent space captures essential features of the input while discarding noise and redundancies. However, it has often been observed that learning with autoen-coders leads to overfitting. To address this challenge, Variational Autoencoders (VAEs)^18 19^ were developed as a probabilistic extension of traditional autoencoders. Although originally conceived for generative modeling, VAEs have also found applications in dimensionality reduction, especially for molecular systems^20^. By imposing a prior distribution, typically a multivariate Gaussian, on the latent space, VAEs introduce regularization into the learning process.^21^ This regularization prevents overfitting by encouraging the latent space to be smooth and structured, leading to better generalization. This modifies the loss function (^ℒ^_*V AE*_) to incorporate both the reconstruction loss and the Kullback-Leibler (KL) divergence term, which regularizes the latent space and is given as

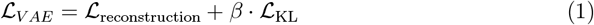

The KL divergence is scaled by the *β* weight, enabling control over the trade-off between reconstruction accuracy and regularization. ^22,23^

A central focus of this work lies in examining the selection of the *β* parameter within the VAE framework. We explore the crucial role, this parameter plays in effectively balancing the competing objectives of accurate data reconstruction and appropriate regularization of the latent space. A higher value of *β* encourages a more regularized and disentangled latent space at the cost of reconstruction quality. Whereas a lower *β* prioritizes accurate reconstruction but may lead to an entangled latent space. A *representation* is considered *disentangled* if it can be broken down into multiple subspaces, each of which aligns with a unique symmetry transformation and can be independently transformed without affecting the others.^24^ Therefore, *β* must be tuned either through visual inspection heuristics or by using disentanglement metrics, such as those obtained by training a simple linear classifier on the latent space of a trained VAE in a supervised manner. However, in the case of MD trajectories, where the data is unsupervised, disentanglement must be assessed through alternative approaches. These include evaluating the clustering of latent representations into meaningful macrostates, analyzing free energy surfaces reconstructed from the latent space, or assessing the ability of the model to capture slow collective motions that are critical to MD studies. While disentanglement focuses on separating independent factors of variation, the latent space in MD studies must also preserve thermodynamic and dynamical relevance, which requires a careful choice of *β* parameter. Metrics such as the Generalized Matrix Rayleigh Quotient (GMRQ) score^25^ and Variational Approach to Markov Processes-2 (VAMP2) score^26^ are essential tools for evaluating the quality of the latent space in terms of its ability to capture slow collective variable, and have been used to monitor the effect of *β* in training VAEs in this work.

We have also examined on how to improve the VAE model’s ability to learn *disentangled* representations. ^27–30^ Early efforts to achieve disentangled representations often relied on prior knowledge of the factors driving data generation.^31–39^ While several fully unsupervised methods have been developed to learn disentangled factors, these approaches have generally struggled to scale effectively to datasets beyond toy datasets. One of the ways to improve disentanglement is by annealing the *β* parameter.^40,41^ While annealing presents an alternative, it does not provide a universal solution. The effectiveness of annealing often depends on the specific dataset and the chosen annealing schedule, and posterior collapse may still occur.^42,43^ Finding the optimal settings can vary significantly across different problems and requires doing expensive parameter sweeps, which can be computationally expensive and time-consuming, especially for high dimensional systems.

In this work, we introduce a novel approach for determining the optimal *β* parameter for training VAEs. Instead of arbitrarily choosing *β* values, we utilize the Fraction of Variance Explained (FVE) score to identify the optimal *β* that maximizes the model’s ability to represent the underlying data distribution and avoid posterior collapse. Our findings have been applied to two systems, Trp-cage and *α*-Synuclein (*α*S). Trp-cage is a 20 residue, fast folding miniprotein, which folds in ≈ 4 *µs*.^44^ *α*S^45^ is an intrinsically disordered protein and is associated with movement disorders such as parkinsons disease, multiple system atrophy and in neurodegenerative disese such as dementia with Lewy bodies. ^46^ Identifying slower collective variables (CVs) is critical for understanding the folding mechanism as in Trp-cage^47^ or capturing the key dynamics of *α*S that drive their functional flexibility. We have trained VAEs with the optimal *β* using three different annealing methods and compared them with the regular VAE model, and found that annealing outperformed their non-annealed conterparts in capturing slower collective variables. Our approach highlights the utility of annealed VAEs in uncovering meaningful latent representations that refine state space discretization, which is critical for subsequent building of Markov State Models (MSM).

## Results

### Probabilistic Approach to Dimensionality Reduction using Variational autoencoder

VAE is composed of two parts, an *encoder* and a *decoder*. The encoder compresses the high dimensional input data *x* into a lower dimensional continuous latent vector *z* which is sampled from a learned approximate posterior distribution. This posterior is regularized to be close to a predefined prior distribution *p*(*z*), typically a standard normal distribution. The job of the decoder is to reconstruct the original data from the conditional distribution *p*_*θ*_(*x*|*z*). Thus the marginal likelihood of *x* is given as

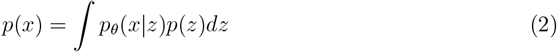

However, the integral is intractable because it involves integrating over all possible values of the latent variable *z* to normalize the posterior distribution, which is given as

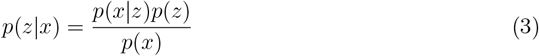

To address this, variational inference is used, in which the true posterior *p*_*θ*_(*z*|*x*) is approximated by a variational distribution *q*_φ_(*z*|*x*), which is parameterized by the encoder using a mean vector and a variance vector. The closeness of the approximated and the true posterior can be quantified using the KL divergence between them and is given as

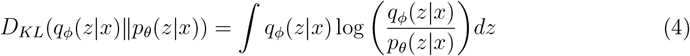

Upon expanding and rearranging, we have

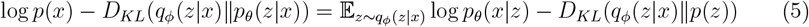

In VAEs, we want to maximize the log likelihood of the true data, log *p*(*x*) and simulataneously minimize the distance between the true and approximated posterior. Thus the training objective becomes minimizing the negative of the left hand side of the above equation. The loss function _*ℒV AE*_ becomes,

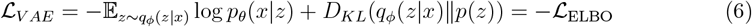

The loss function is the negation of Evidence Lower Bound (ELBO). Minimizing the loss is equivalent to maximizing the ELBO, which serves as a lower bound to log *p*(*x*). This encourages accurate reconstructions while enforcing a structured latent space which aligns the approximate posterior to the prior.

### Learning disentangled representation using *β*-VAE

In many cases, a standard VAE does not ensure that the latent variables capture independent features of the data. This limitation led to the development of *β*-VAE, which explicitly aims to disentangle the factors of variation in the data. This is achieved by constraining the distance between the approximate posterior and the prior to remain below a small constant, δ,

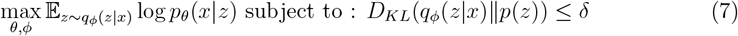

To handle the constraint, the Lagrangian is defined as

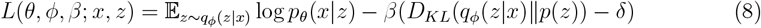

where *β* is the lagrange multiplier. Using the complementary slackness condition from the Karush-Kuhn-Tucker (KKT) conditions, the necessary conditions for optimality in constrained optimization problems can be derived. Since δ is fixed, −*β*δ is simply a constant offset in the Lagrangian. Constant terms do not affect the gradient with respect to the model parameters, so we can safely omit −*β*δ in the final objective. Consequently, the loss function of the VAE^48^ reduces to

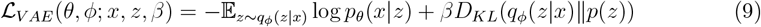

### Augmenting disentanglement through annealing of the *β* parameter

The choice of *β* parameter plays a critical role in balancing the trade-off between reconstruction quality and the regularization of the latent space. A higher value of *β* encourages a more regularized and disentangled latent space at the cost of reconstruction quality, whereas a lower *β* prioritizes accurate reconstruction but may lead to an entangled latent space. In dimension reduction, the primary objective is to preserve as much information from the original data as possible, which can be accomplished by using a lower value of *β*. However this may lead the approximate posterior to collapse to the prior, a phenomenon known as KL vanishing problem. To better understand the impact of the *β* parameter and its role in balancing these trade-offs, we examined the *D*_*KL*_ term,^49–52^ by decomposing it into two components:

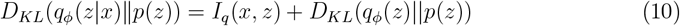

The mutual information (MI) term *I*_*q*_(*x, z*) measures the degree to which the encoded representation *z* captures the unique features of the input data *x*, so that the decoder can faithfully reconstruct it. A higher MI enhances the correlation between the latent variables and the data, promoting a reduction in the extent of KL vanishing. The marginal KL term, *D*_*KL*_(*q*_φ_(*z*)‖ *p*(*z*)), evaluates how well the aggregated posterior, *q*_φ_(*z*), aligns with the prior distribution. Combining the above two equations, the loss function can be written as

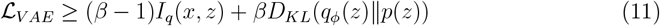

In case of a standard VAE, where *β* = 1, the first term disappears, leaving only the KLD term. Training the model focusses solely on aligning the aggregated posterior with the prior with no incentive to maximize the MI, which leads to posterior collapse. As a result the latent space captures no meaningful information about the input data. However annealing of the *β* parameter^40,41^ can alleviate such problems thereby providing a better latent representation, which retains the best of both worlds. To begin with, we start with a low value of *β* which allows the model to capture the underlying pattern in the data more freely without emphasising much on the prior *i*.*e*. the model emphasizes on maximizing the mutual information. This creates a latent representation that is meaningful but not regularized. As the training progresses, the increase in the *β* value makes the latent space more regularized thereby aligning it to the prior distribution, rather than solely focussing on reconstructing the input data. When *β* reaches 1, the weighted loss function is equivalent to the true variational lower bound. This gradual approach of annealing *β*, enables the model to alleviate overlapping or entangled features, thereby improving the model’s ability to learn disentangled factors of the latent space.

### Our proposed protocol for optimizing *β* with FVE

While annealing *β* to 1 improves the model’s ability to balance reconstruction and regularization, in practice, this may not always yield optimal results. One way to improve is to cyclically anneal the *β* value to 1.^41^ But it requires selection on the number of cycles and also needs to be trained for large number of epochs (so that the annealing rate is slow and reduce the risk of posterior collapse), thereby increasing training time. In this work, we give an alternate way in which we anneal the *β* parameter in just one annealing cycle. Instead of annealing *β* to 1, we anneal it to a lower value. Literature survey^53,54^ indicates that, the *β* parameter have been arbitrarily set during the training of standard VAE models. However, this is ineffective as the results may change upon choosing a different *β* value. We have, instead used FVE score^55^ as a metric to find an optimal value of *β* for training the model, which is given as

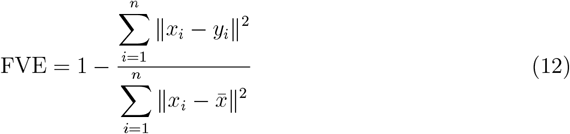

where 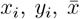 and *n* represent the input, output, mean of the input and total number of features respectively. The FVE score is calculated after each training epoch, and we select the *β* value corresponding to the highest value of FVE score in the full training cycle. We chose this value as this score suggests that the model at that specific *β*, has been able to retain the maximum variance of the data during reconstruction. In this work, the rate of this increase is tuned using three different schedules, namely linear^56,57^(equation 13), cosine^58^(equation 14) and logistic^40^(equation 15) as defined by the respective equation

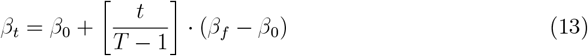

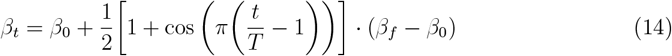

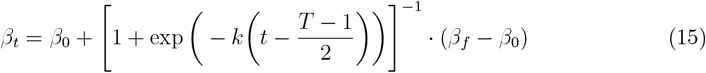

where *t* is the current epoch, *T* is the total number of epochs, *β*_0_ is the starting value of *β, β*_*t*_ is the *β* at epoch *t* and *β*_*f*_ is the final value of *β. k* determines the steepness of the logistic curve and is set to 10/(*T* − 1).

### Training of VAEs using our method

The training protocol begins with using the C_*α*_ pairwise distance matrix as the input feature vector for the model. The VAE is then trained using different annealing schemes, with the *β* parameter gradually annealed to a value of 1. As a control, a standard VAE model is also trained with *β* fixed to 1. During training, the FVE score is monitored as a function of *β*, and the optimal *β* value is determined as the one corresponding to the maximum FVE score. Once the optimal *β* values are identified, the VAE is retrained using each annealing scheme at its respective optimal *β*. The best-performing annealed VAE model is selected based on the highest VAMP2 score, calculated from the mean vector and compared with the non-annealed counterpart (see Figure 1(d)). The mean vector from the selected model is then used to construct MSM.

**Figure 1.**
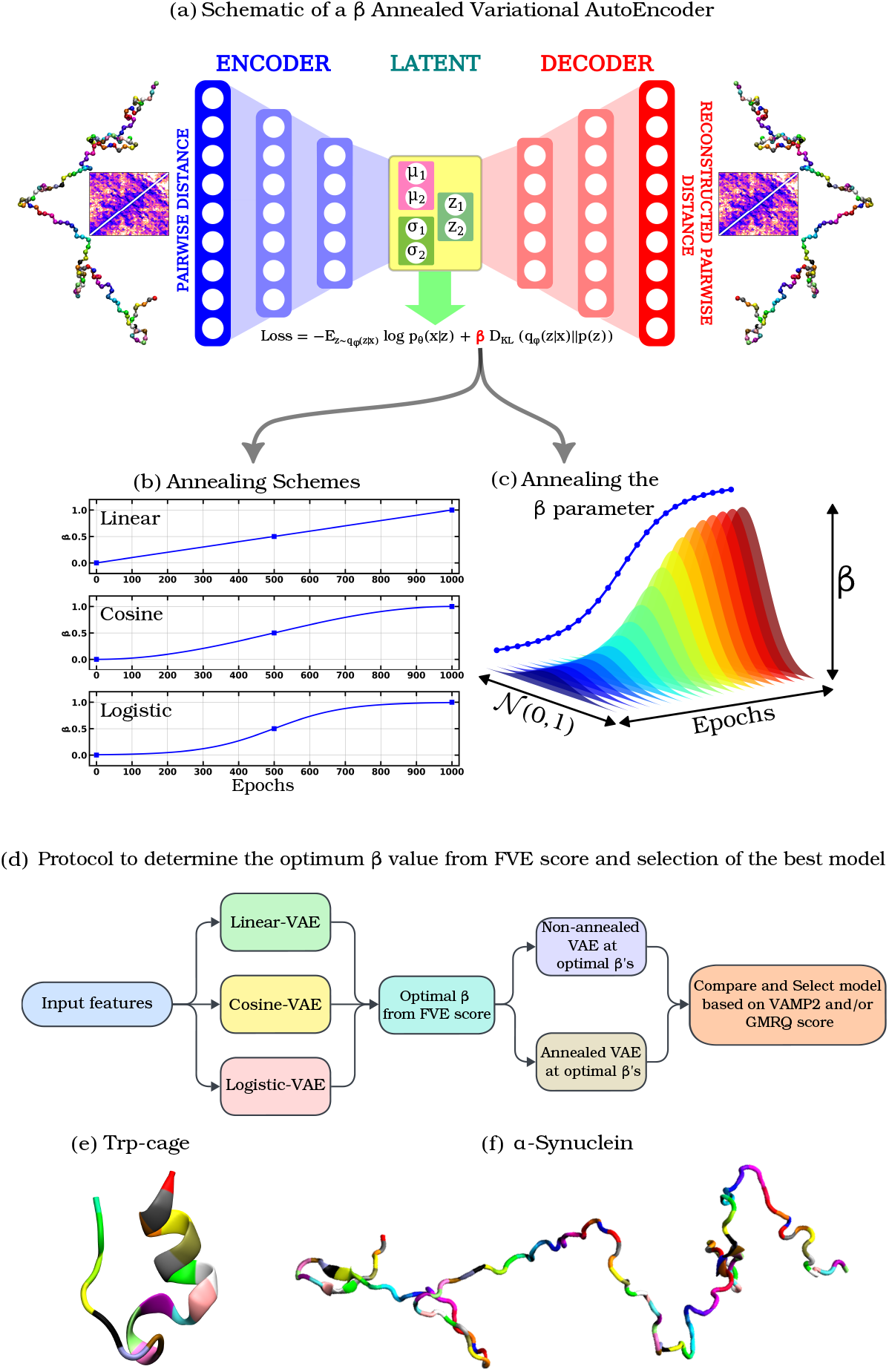
(a) Schematic of a *β* Annealed Variational AutoEncoder (b) Linear, Cosine and Logistic annealing schedules (c) Annealing of the *β* parameter. (d) Protocol to determine the optimum *β* value from FVE score and selection of the best model to construct MSM. Cartoon snapshot of the Trp-cage mini protein and (f) *α*S protein. The different colours correspond to different residues.

## Discussion

### Optimizing VAE for Improved Representation of Protein Free Energy Landscapes

We initially trained a standard VAE model (*β* fixed to 1) and annealed VAE model (*β* annealed to 1) for the Trp-cage protein (refer to ‘Method (Overview of training data) section’). The mean vector produced by the encoder in the latent space of the trained model was used to construct the free energy surface (FES) and shown in Figure 2, for both the standard VAE and annealed VAE across all annealing schemes. The FES revealed a single minimum for all models, indicative of posterior collapse, where the model fails to capture distinct states. Notably, the standard VAE exhibited a more pronounced and collapsed minimum compared to the annealed VAE. This suggests that while the annealing process partially mitigates posterior collapse, it ultimately fails to avoid it, indicating room for further improvement in balancing reconstruction and regularization.

**Figure 2.**
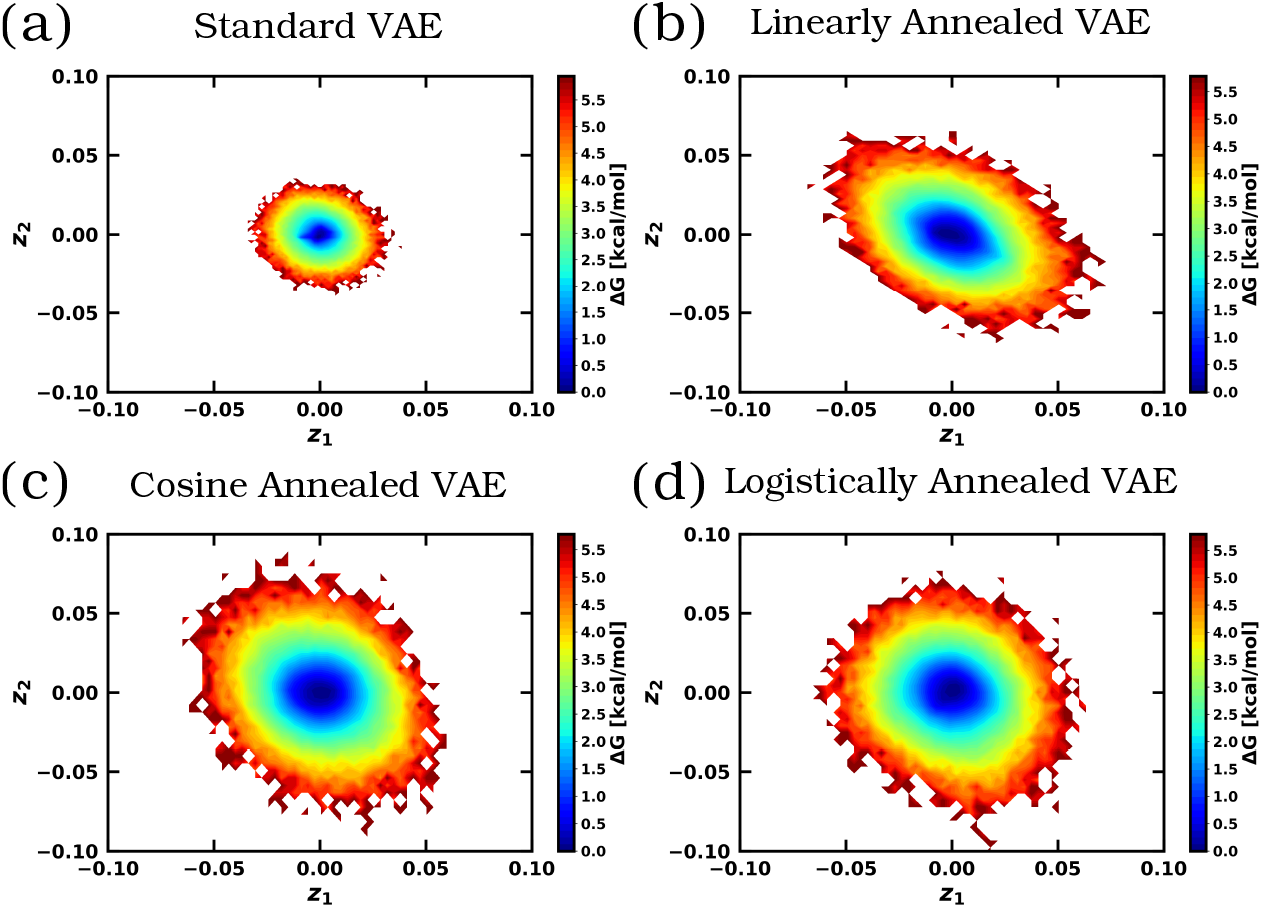
FES of Trp-cage using the latent space of (a) Standard VAE (b) Linearly annealed VAE (c) Cosine annealed VAE and (d) Logistically annealed VAE. *β* is annealed to 1.

To investigate this collapse, we have monitored the FVE score over the entire training cycle as shown in Figure 3(a). The initial negative FVE score reflects that the model’s reconstruction is poor at the start of training. This is expected, as the VAE starts with randomly initialized parameters, and the latent space is not yet structured in a way that effectively captures the data variance. As the training progresses, the model learns to reconstruct the data more effectively, leading to an increase in the FVE score. The peak corresponds to the point where the model has reached an optimal balance between the variance and regularization. After this point, as *β* continues to increase, the weight of the KL divergence term grows, leading to stronger regularization of the latent space. This regularization forces the latent variables to conform more closely to the prior distribution. Consequently, the ability of the model to explain the variance decreases, resulting in a decline in the FVE score. Eventually, as *β* reaches to 1, the regularization fully dominates, and the model prioritizes aligning the latent space with the prior, over capturing meaningful variance from the data. At this stage, the FVE score flattens out to near zero values, indicating that the latent space is no longer effectively representing the data. The latent variables contribute no useful information, and the decoder essentially defaults to reconstructing the mean of the data. In comparison, the standard VAE is trained with *β*=1 from the begining, which forces the model to balance reconstruction and regularization throughout training. This early regularization restricts the model’s capacity to capture meaningful variance, leading to a more severe posterior collapse compared to the annealed VAE.

**Figure 3.**
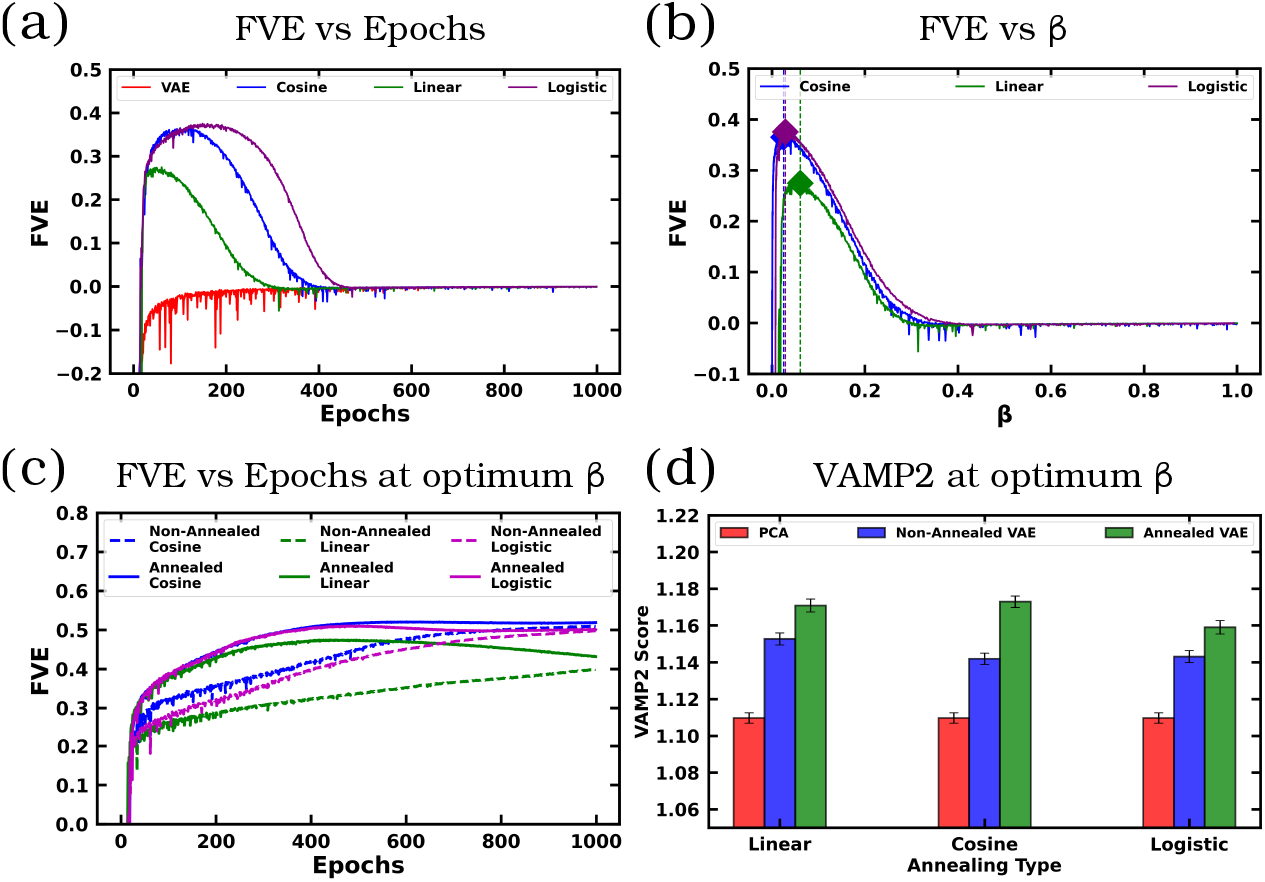
(a) Variation of FVE value with respect to epochs for VAE and the annealing schemes. Change in FVE as a function of *β* for different annealing schemes. The optimum value of *β* is labelled by the diamond marker which is 0.061, 0.025 and 0.029 for linear, cosine and logistic annealing respectively. (c) Variation of FVE values with respect to epochs for models trained at their optimum *β* values. (d) Comparison of the VAMP2 score between PCA, non-annealed VAE and annealed VAE for different annealing schemes for lag time of 10 ns.

To address posterior collapse, we retrained the VAE model using the optimal *β* value corresponding to the highest FVE score achieved by the annealed VAE across all annealing schemes. Similarly, each VAE model was trained by annealing the *β* to its respective optimal *β* value (see Figure 3(b)). We evaluated six models in total: three from the non-annealed VAE and three from the annealed VAE. We analyzed the FVE scores of the non-annealed VAE and annealed VAE and found that while the FVE scores for both models eventually plateau to a similar value by the end of training, their trajectories differ significantly. The annealed VAE demonstrates a rapid increase in the FVE score during the initial stages of the training before plateauing. In contrast, the non-annealed VAE exhibits a slower, more gradual increase in the FVE score, reflecting its difficulty in capturing the variance and lags behind the annealed VAE during the entire training cycle (see Figure 3(c)). While both models eventually converge to a similar FVE score, the annealed VAE’s training trajectory suggests it is more efficient at capturing meaningful data variance early on, mitigating posterior collapse, and structuring the latent space in a way that may be more useful for down-stream tasks. To evaluate the effectiveness of the two models, we compared their VAMP2 scores, which measure the ability of the latent space to capture slow collective variables, and compared it with the top two principal components from PCA. Our results demonstrate that the five-fold cross-validated VAMP2 scores for the annealed VAE consistently outperformed those of the non-annealed VAE and PCA across all annealing schemes (see Figure 3(d)). This highlights that annealing the *β* parameter significantly enhances the model’s ability to capture slow collective variables compared to both non-annealed VAEs and PCA.

To further assess the quality of the latent space, we analyzed its disentanglement by clustering the FES using a Gaussian Mixture Model (GMM) (see Figure 4(a)-(b) and Figure S1). The optimal number of clusters was determined by evaluating Bayesian Information Criterion (BIC) scores across different cluster counts (see Figure 4(c) and Figure S2). The slope of the BIC curve indicated that four clusters provided the best trade-off between model complexity and goodness of fit for different annealing schemes. After clustering, we assessed the quality of the identified clusters using the silhouette score, which quantifies how well a data point fits within its assigned cluster compared to others. Higher silhouette scores indicate better-defined and more distinct clusters. Our analysis revealed that the silhouette scores for the four clusters derived from the annealed VAE were consistently higher across all annealing schemes compared to the non-annealed VAE (see Figure 4(d)). This suggests that annealing the *β* parameter enhances the disentanglement of representations in the latent space, resulting in more distinct and interpretable clusters. The points corresponding to each cluster were color-coded, highlighting the distinct regions of the latent space occupied by the four clusters. In the case of the non-annealed VAE, the clusters appeared less well-separated, reflecting the challenges in disentangling representations without annealing. By contrast, the annealed VAE exhibited a more distinct and organized clustering in the latent space, underscoring the improved separation and representation achieved through annealing (see Figure 4(e)-(f)), as further supported by the consistently higher silhouette scores (see Figure 4(c)-(d) and Figure S3).

**Figure 4.**
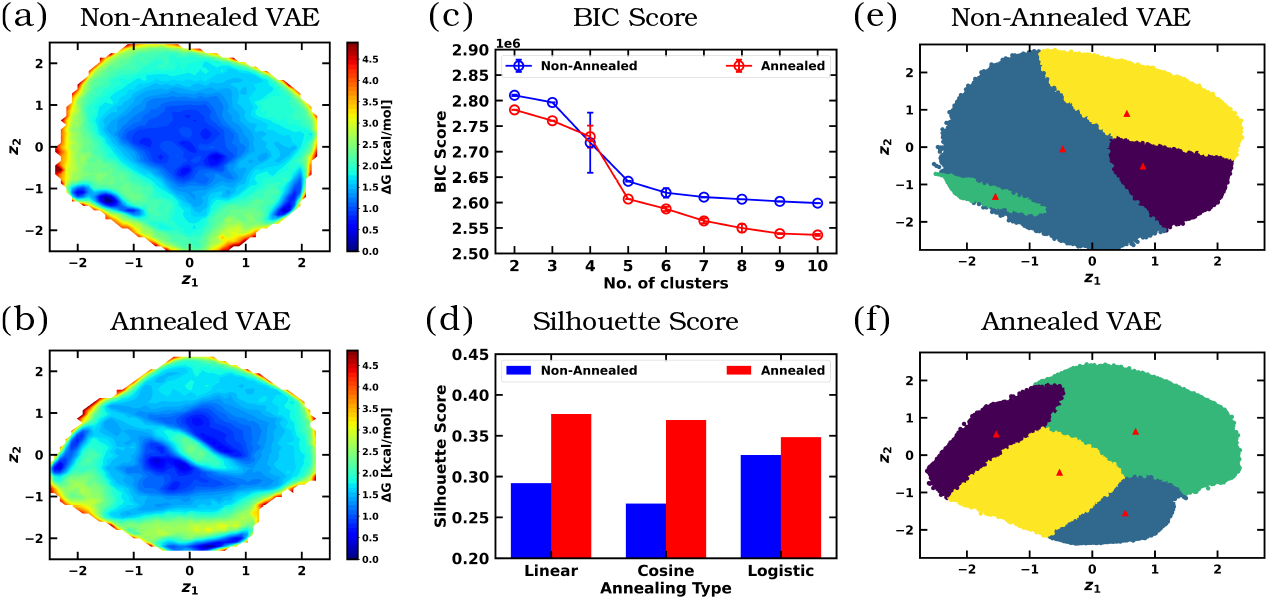
FES of Trp-cage using the latent space of (a) Non-annealed cosine VAE (b) Cosine annealed VAE (c) BIC score as a function of GMM clusters for the Non-annealed cosine and Cosine annealed VAE (d) Sihouette score as a function of GMM clusters for all the annealing schemes. The clusters identified by the GMM from the latent space of (e) Non-annealed Cosine VAE and (f) Cosine annealed VAE. The cluster centers are marked in red triangles.

### Refining Kinetic and Structural Understanding of *α*S with Annealed VAEs

We then investigated the effect of annealing on a more complex system, *α*S (refer to ‘Method (Overview of training data) section’). To achieve this, we identified the optimal *β* parameter and trained both non-annealed and annealed VAE models using this value (see Figure S4). We then compared the VAMP2 scores derived from the latent space of the models. Consistent with previous observations, the annealed VAE outperformed the non-annealed VAE, achieving higher VAMP2 scores across all annealing schemes (see Figure S5). The FES constructed from the latent space of the annealed VAE demonstrated a more dispersed distribution of latent representations compared to the non-annealed VAE alongwith an increased number of local minima, a characteristic feature of IDPs (see Figure 5(a)-(b) and Figure S6). To better understand the dynamic transitions between the metastable states and gain insights into the underlying kinetics, we performed kinetic modeling by constructing a Markov State Model (MSM). We used k-means clustering to discretize the latent space into microstates, with the number of clusters ranging from 50 to 1000. To determine the optimal number of microstates, we employed both the GMRQ score and VAMP2 score as metrics. In both of these metrics, the annealed VAE outperformed the non-annealed VAE (see Figure 5(c) and Figure S7-S8), indicating that the annealing process enhanced the quality of the latent space representation. We chose 500 microstates to ensure a sufficiently detailed representation of the system’s dynamics. A transition matrix was then built by counting the number of transitions among the microstates at lag times ranging from 1 ns to 200 ns. Using this matrix, we computed the implied timescales (ITS) for both the non-annealed and annealed VAE (see Figure 5(e)-(f)). The ITS values for the annealed VAE were higher than those for the non-annealed VAE, indicating that the annealed VAE captures slower relaxation processes more effectively. This suggests that the annealing process enhances the model’s ability to identify transitions between more kinetically distinct or stable states. Similar trends were observed across other annealing schemes as well (see Figure S9).

**Figure 5.**
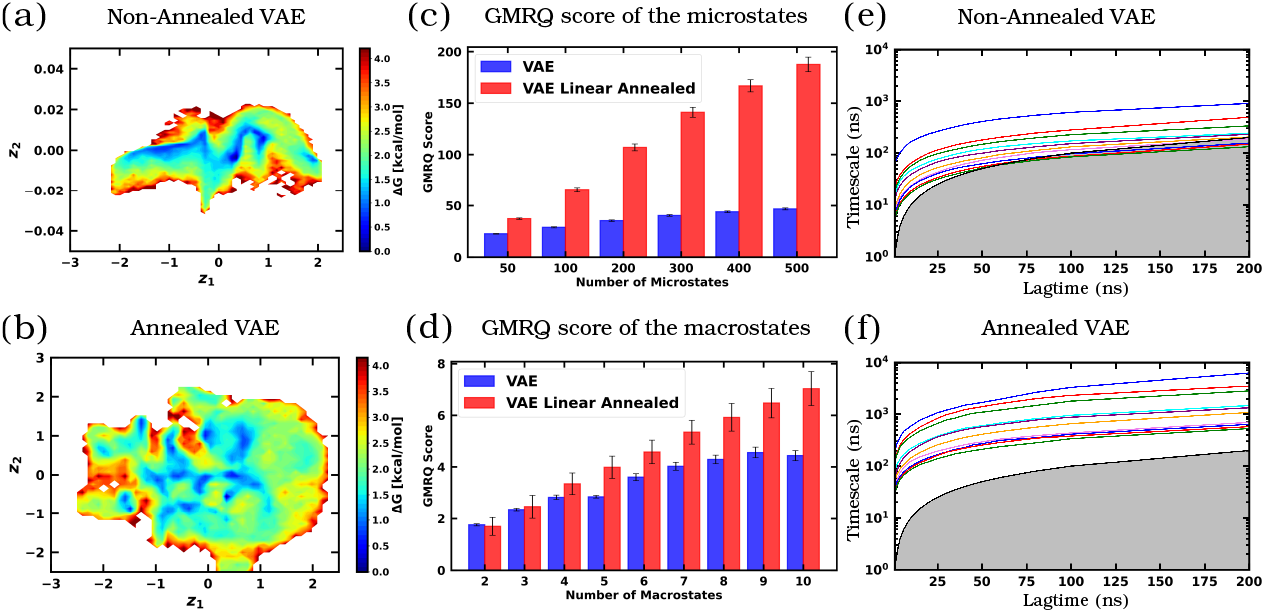
FES of *α*S using the latent space of (a) Non-annealed linear VAE (b) Linear annealed VAE. GMRQ scores of the Non-annealed linear VAE and Linear annealed VAE for different number of (c) microstates and (d) macrostates. ITS of the (e) Non-annealed linear VAE and (f) Linear annealed VAE.

To coarse-grain the microstates, Perron Cluster Cluster Analysis (PCCA+) was performed at a lag time of 100 ns. The number of macrostates was varied between 2 and 10. The five-fold cross-validated GMRQ scores from the training dataset indicate that, regardless of the number of macrostates, the annealed VAE consistently achieves higher GMRQ scores across all annealing schemes (see Figure 5(d) and Figure S7). Finally, we discretized the MSM for all the annealing schemes into six macrostates (see Figure 6 and Figure S10) and Chapman-Kolmogorov (CK) test was performed to verify the Markovianity of the model for all non-annealed and annealed VAEs (see Figure S11-S13). The CK test showed that the linear and cosine annealed VAEs outperformed their non-annealed counterparts, with prediction and re-estimation in good agreement. However, the logistically annealed VAE performed slightly worse than the non-annealed case. For the linearly annealed VAE, the macrostates were characterized using inter-residue contact maps. These contact maps revealed distinct structural features for each state, confirming their uniqueness and highlighting the ability of annealed VAE to capture structurally significant conformations of the system (see Figure 7(a)-(f)).

**Figure 6.**
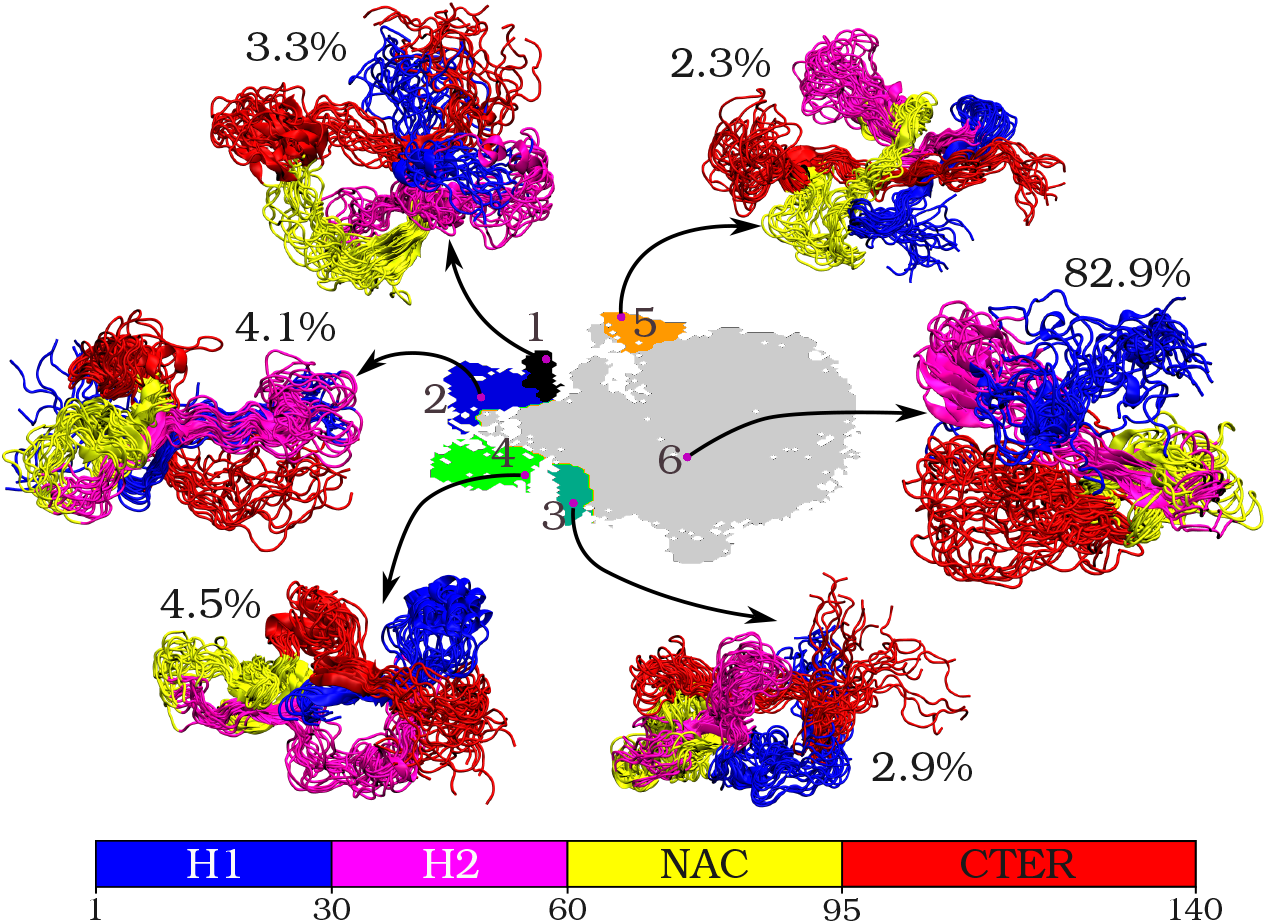
Coarse grained MSM macrostates of *α*S using the latent space of Linear annealed VAE, with their representative structures and their population. The bar below represents different regions of *α*S.

**Figure 7.**
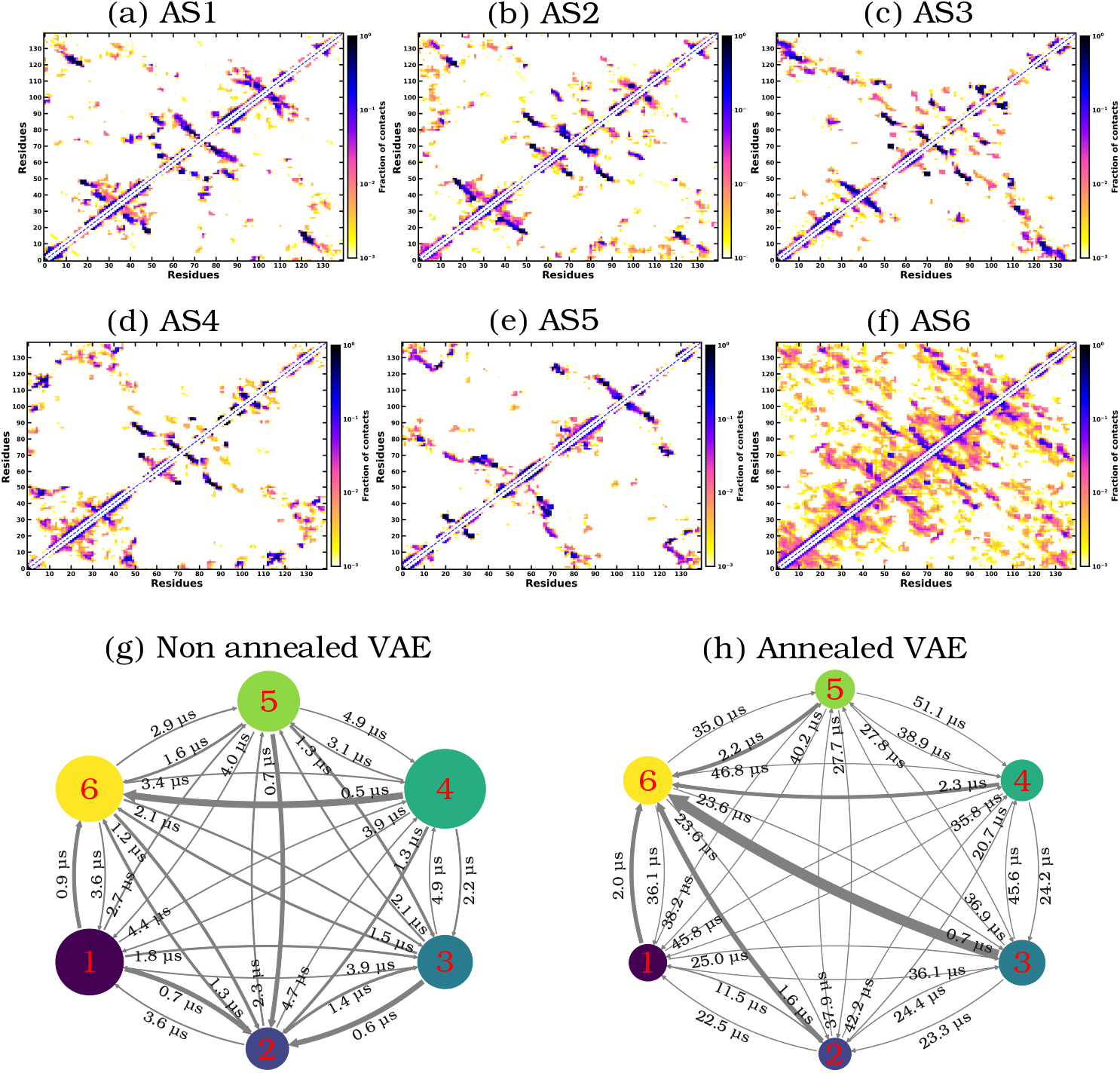
(a)-(f) Ensemble averaged backbone contact map for the six Linearly annealed VAE states AS1-AS6. MFPT among the macrostates of (g) Non-annealed linear VAE and (h) Linearly annealed VAE.

The residue-wise contact map for the linearly annealed states of *α*S highlights significant interactions across its four structural regions. The N-terminal region, characterized by a highly conserved hexamer motif (KTKEGV), exhibits a tendency to form helices and can be divided into two helical segments: H1 (residues 1-30) and H2 (residues 31-60) when associated with micelles. This is followed by a central hydrophobic nonamyloid-*β* component (NAC) region (residues 61 to 95) and C terminal region (residues 96 to 140). The different annealed states (AS) obtained from the MSM macrostate, reveal distinctive structural motifs and inter-domain interactions within *α*S. In AS1, notable *β*-sheet contacts are observed between ALA17-GLU28 in the H1 region and VAL37-VAL49 spanning the H2 region, while the NAC region forms *β* sheets with LYS60-ALA69 interacting with ALA78-SER87, emphasizing the structural organization in this aggregation-prone area. Transitioning to AS2, a *β*-sheet between VAL49-ALA56 (H2) and GLU83-ALA90 (NAC) links these regions, with MET5-ALA17 (H1) and PRO120-ASP135 (CTER) forming another *β* sheet, highlighting inter-domain interactions and structural connectivity across the regions. In AS3, significant *β*-sheet contacts between ALA30-TYR39 (H2) and PRO108-LEU113 (CTER) suggest long range connectivity, while MET1-ALA17 (H1) and ASP115-ASP135 (CTER) indicate long-range NTER-CTER interactions. The AS4 state reveals an antiparallel *β*-sheet between ASP4-ALA17 (H1) and LYS45-HIS50 (H2), alongside long-range contacts between ALA19-VAL40 (H1) and ALA124-GLY132 (CTER), suggesting a well-organized structural alignment across multiple domains. In AS5, inter-region interactions are highlighted by contacts between MET5-LYS43 (H1) and THR64-VAL82 (NAC), as well as *β* sheets between ALA90-LEU100 (NAC) and GLY106-VAL118 (CTER), which contribute to the stability of the structure. Extended *β*-sheet arrangements are also seen between MET5-GLY14 (H1) and GLU123-TYR133 (CTER), linking distant regions of *α*S. Finally, the AS6 state, characterized by a large population, demonstrates widespread inter-region connectivity, with notable interactions between GLY68-ALA78 (NAC) and GLU123-GLU137 (CTER), as well as GLU20-LYS32 (H1) and TYR125-TYR136 (CTER), further illustrating the complex and dynamic network of interactions that govern the conformational behavior of *α*S across different annealed states (see Figure 7(a)-(f)).

We have further calculated the Mean First Passage Time (MFPT) between the metastable states for both the non-annealed VAE and the annealed VAE. Our analysis reveals that the timescales reported by the non-annealed VAE are an order of magnitude lower compared to those from the annealed VAE. This significant discrepancy in timescales indicates that the non-annealed VAE latent space over-simplifies the system’s dynamical landscape resulting in overestimated rates of conformational changes. Consequently, the system’s kinetics seems distorted where transitions between states appear unrealistically fast (see Figure 7(g)-(h)).

This study demonstrates the efficacy of annealed VAEs in capturing the complex conformational landscapes of proteins, both for a model Trp-cage protein and an intrinsically disordered system, *α*S. By systematically selecting the annealing parameter *β* through the FVE score we were able to mitigate posterior collapse and ensure the generation of meaningful latent representations. For both Trp-cage and *α*S, annealed VAEs outperformed their non-annealed counterparts in various metrics, including VAMP2 and GMRQ scores, FES analysis, and Markov State Model analysis. The slower convergence velocities and the presence of more distinct minima in the FES for annealed VAEs highlight their ability to capture finer details of the conformational space. The annealed VAE, with its higher reported timescales, captures a more faithful representation of the long-lived states and slow transitions, which are often critical for understanding system behavior. Overall, this work provides compelling evidence that annealed VAEs offer a more robust framework for exploring the conformational dynamics of proteins. The interplay between *β* annealing schedules, the latent space structure, and the performance metrics across both Trp-cage and *α*S systems underscores the importance of optimizing these parameters to capture meaningful and interpretable biological insights.

## Methods

### Neural Network Architecture

To represent protein structures, we utilized C_*α*_-C_*α*_ pairwise distances as input features. For Trp-cage, distances were calculated using all C_*α*_ atoms, whereas for *α*S, distances were computed by skipping every third residue. The neural network architecture for Trp-cage consisted of layers with 190, 128, 64, 32, and 16 neurons, while for *α*S, it included layers with 1081, 512, 256, 64, and 12 neurons. In both cases, the latent space was represented by 2 neurons. The models were trained using stochastic gradient descent (SGD) as the optimizer with a learning rate of 1 *×* 10^−3^ and mean squared error (MSE) was used as the loss function, weighted by a factor of 10. A batch size of 5000 and 100 was used for Trp-cage and *α*S respectively. Hyperbolic tangent (tanh) activation was used in hidden layers, while the output layer utilized a sigmoid activation. Weights were initialized using the *glorot uniform* method. The training was performed for 1000 epochs, with 90% of the data allocated for training and 10% for validation. A schematic of the architecture is shown in Figure 1(a)-(c). The implementation of the *β* annealed VAE is made publicly available in the Github page (https://github.com/subinoyadhikari/beta_annealed_VAE)

### Overview of training data

The training of VAEs was carried out using high-quality, long-timescale molecular dynamics simulation trajectories of two distinct protein systems: Trp-cage and *α*-Synuclein. Specifically, a 100 *µ*s unbiased simulation trajectory of Trp-cage and a 73 *µ*s simulation trajectory of *α*-Synuclein were utilized, both provided by D. E. Shaw Research.^59,60^ These datasets are well-suited for exploring protein dynamics due to their exceptional temporal resolution and extensive sampling of conformational space. The representative structures are shown in Figure 1-(e) and (f) respectively.

## Data and code availability

All data supporting the findings are included within the manuscript. Additionally, the code for training the model can be accessed on GitHub via the following link: https://github.com/subinoyadhikari/beta_annealed_VAE

## Acknowledgments

We acknowledge support of the Department of Atomic Energy, Government of India, under Project Identification No. RTI 4007. We sincerely acknowledge Tata Institute of Fundamental Research Hyderabad, India for providing the support of computing resources. We thank to D. E. Shaw Research for providing us the long MD simulation trajectories of Trp-cage and *α*-Synuclein.^59,60^ JM acknowledges Core Research grants provided by the Department of Science and Technology (DST) of India (CRG/2023/001426).

