## Supplemental texts and figures and tables for "Elucidating Protein Dynamics through the Optimal Annealing of Variational Autoencoders"

### Supporting Information for Elucidating Protein Dynamics through the Optimal Annealing of Variational Autoencoders

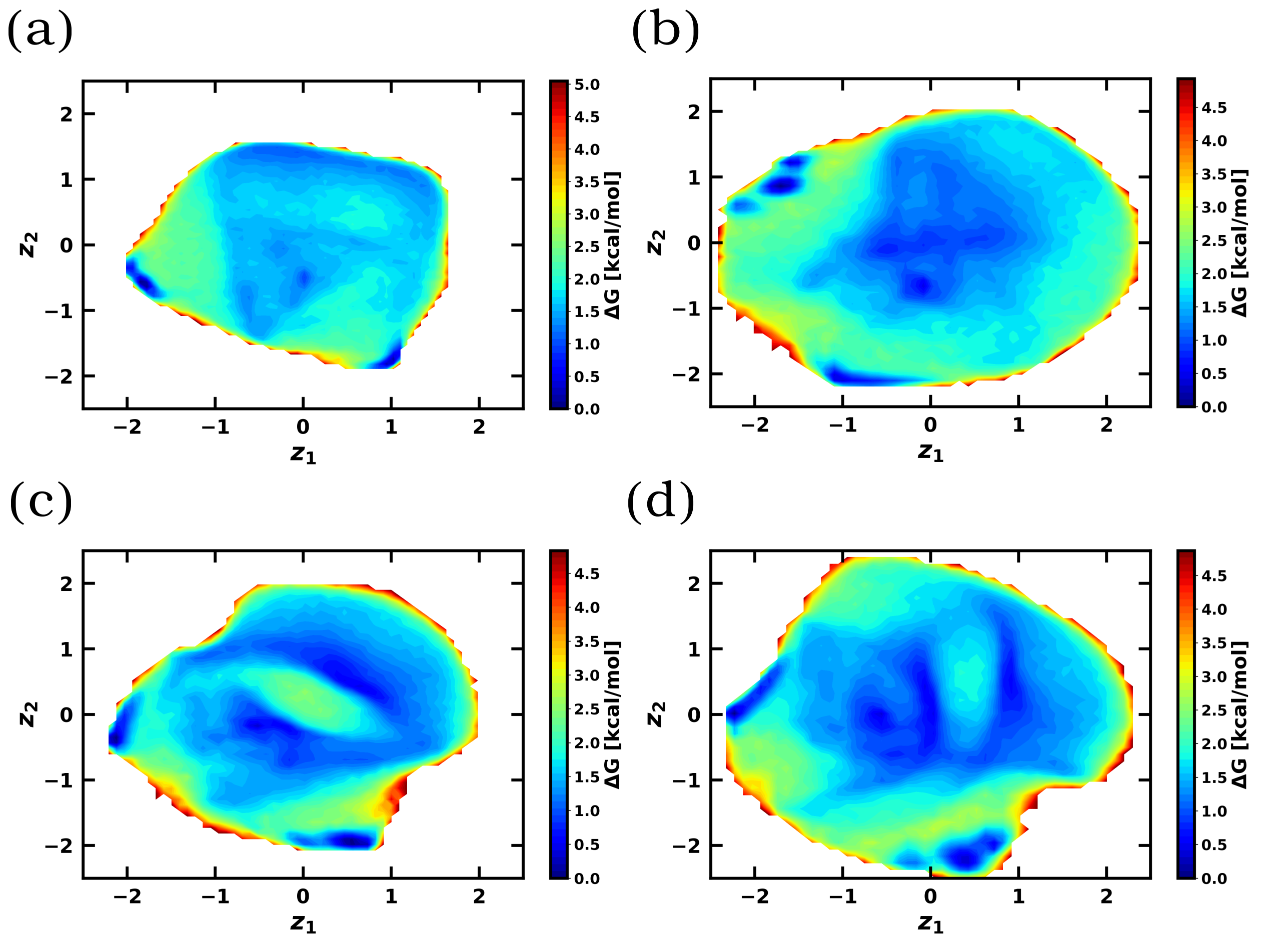


**Figure S1:** Free Energy Surface of Trp-cage for (a) Non-annealed linear VAE (b) Non-annealed logistic VAE (c) Linear annealed VAE and (d) Logistic annealed VAE.

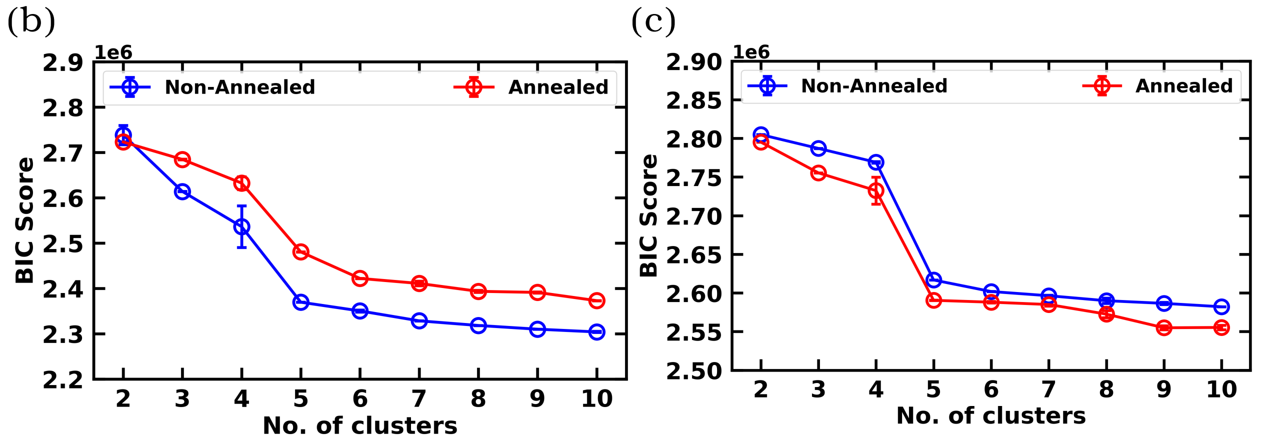


**Figure S2:** BIC score of Trp-cage for (a) Non-annealed linear and annealed linear VAE (b) Non-annealed logistic and annealed logistic VAE.

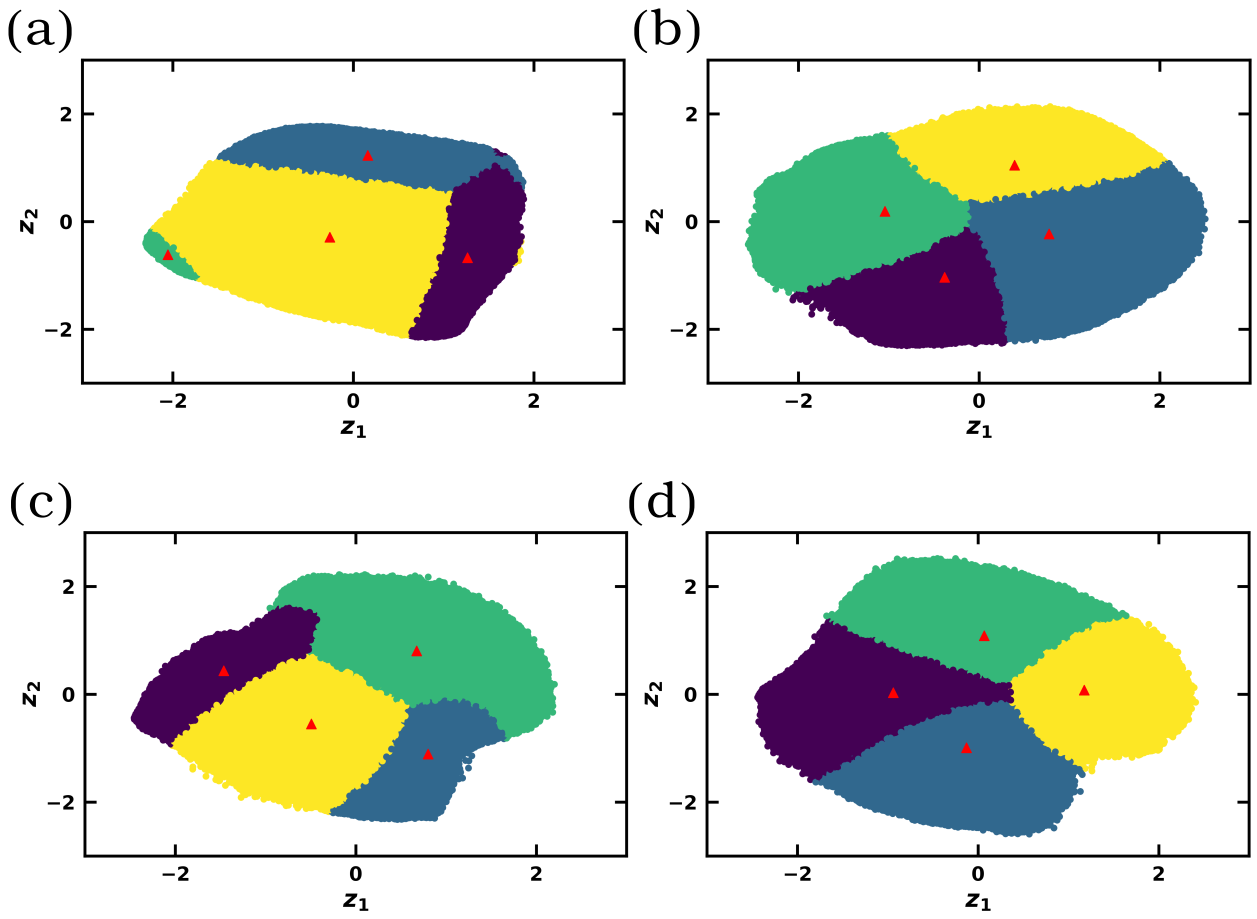


**Figure S3:** GMM cluster of Trp-cage for (a) Non-annealed linear VAE (b) Non-annealed logistic VAE (c) Linear annealed VAE and (d) Logistic annealed VAE.

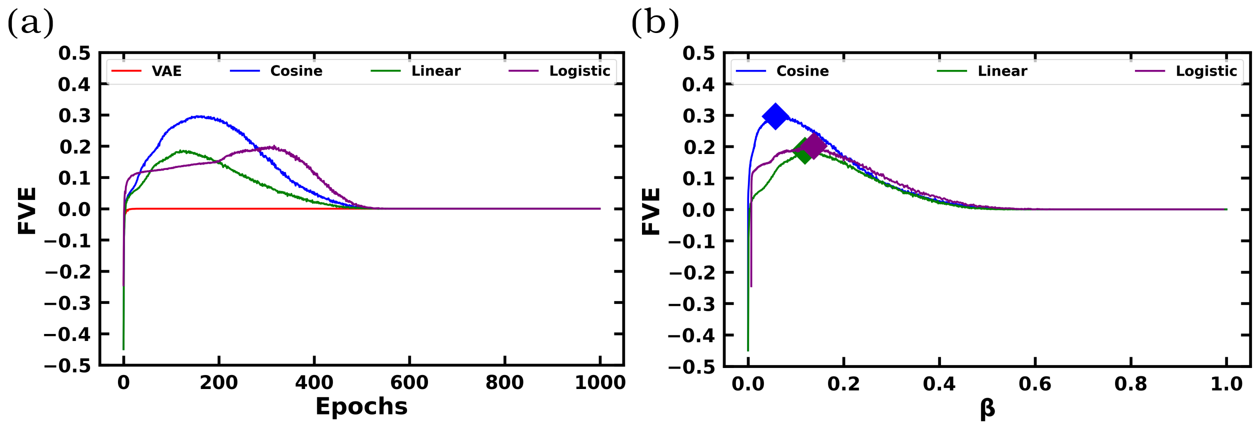


**Figure S4:** (a) Variation of FVE of $\alpha$S with respect to Epochs (b) Variation of FVE of $\alpha$S with respect to $\beta$.

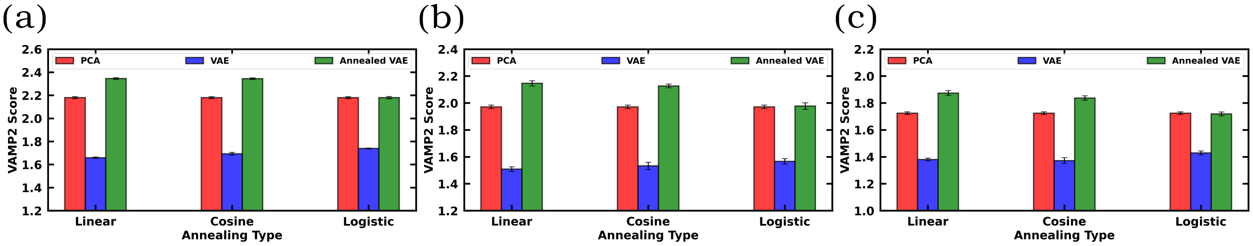


**Figure S5:** VAMP2 Scores of $\alpha$S for PCA, non-annealed VAE and annealed VAE for lag times of (a) 10 ns (b) 20 ns and (c) 50 ns.

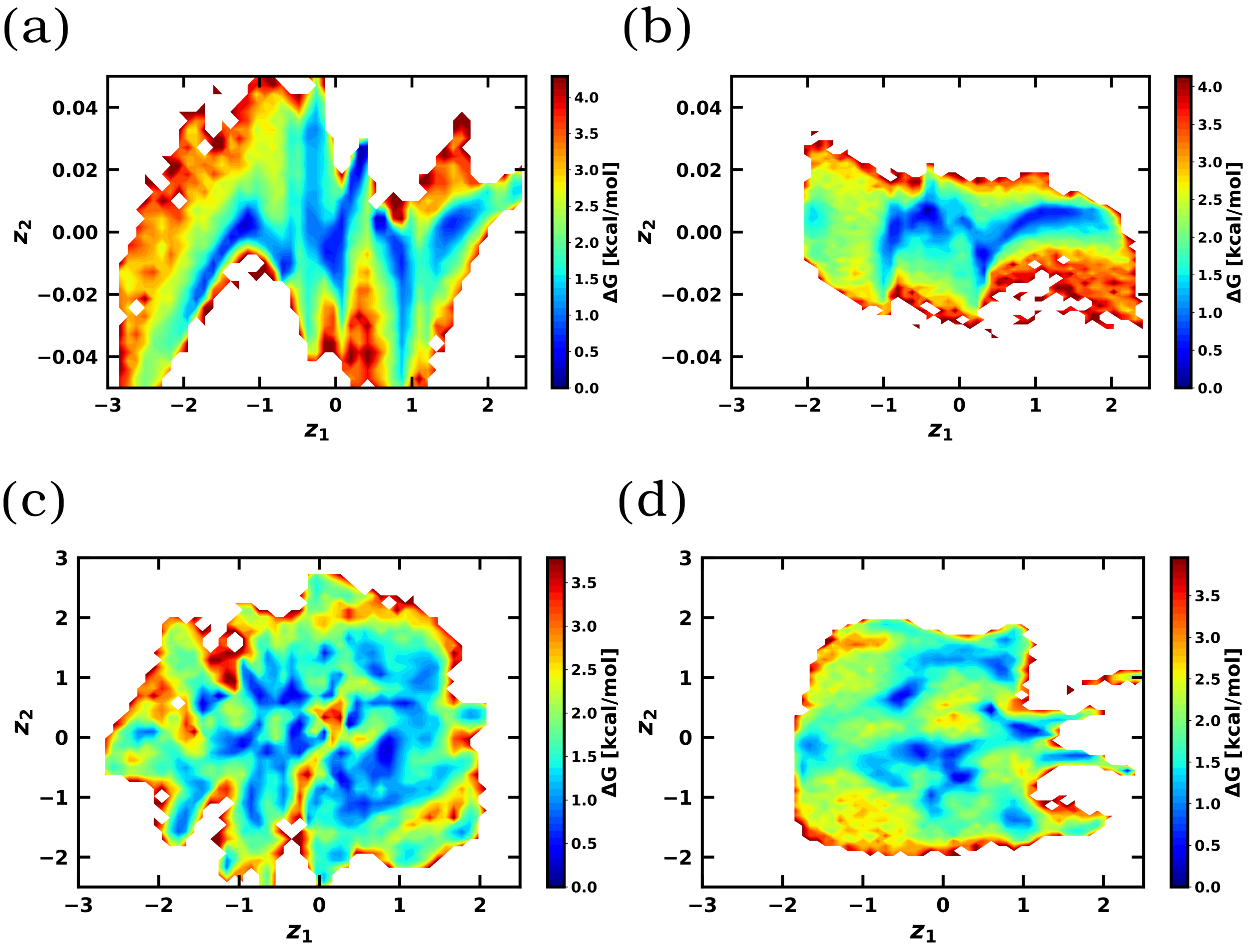


**Figure S6:** Free Energy Surface of $\alpha$S for (a) Non-annealed cosine VAE (b) Non-annealed logistic VAE (c) Cosine annealed VAE and (d) Logistic annealed VAE.

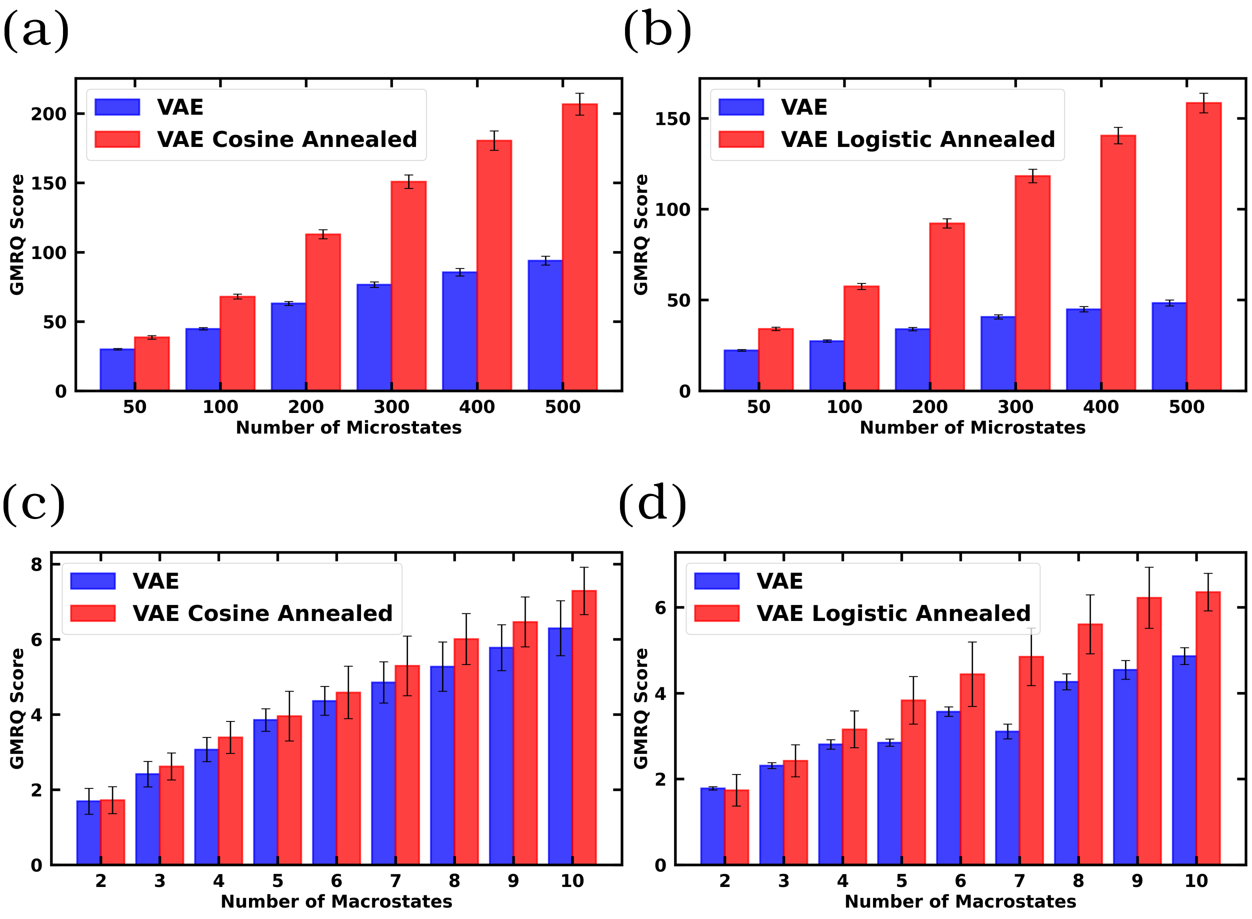


**Figure S7:** GMRQ scores of $\alpha$S for (a) Non-annealed cosine and Cosine annealed VAE for different number of micostates (b) Non-annealed logistic and Logistic annealed VAE for different number of microstates. (c) Non-annealed cosine and Cosine annealed VAE for different number of macrostates (d) Non-annealed logistic and Logistic annealed VAE for different number of macrostates.

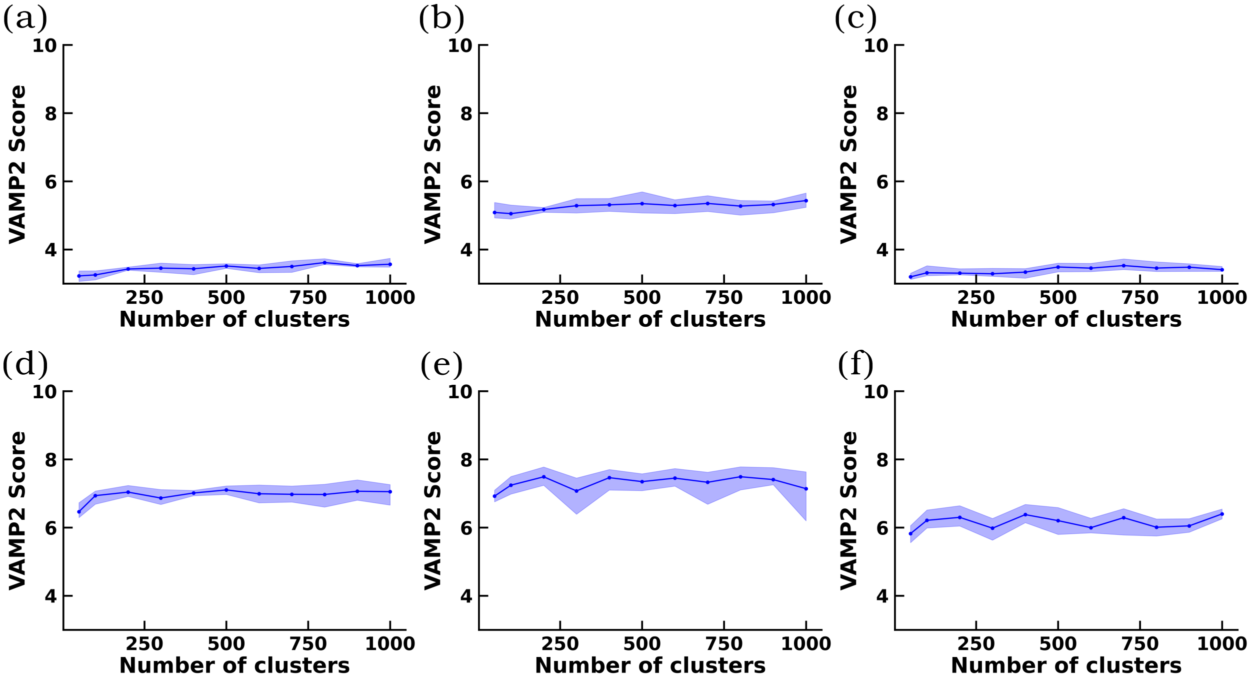


**Figure S8:** VAMP2 Score of $\alpha$S for (a) Non-annealed linear VAE (b) Non-annealed cosine VAE (c) Non-annealed logistic VAE (d) Linear annealed VAE (e) Cosine annealed VAE and (f) Logistic annealed VAE.

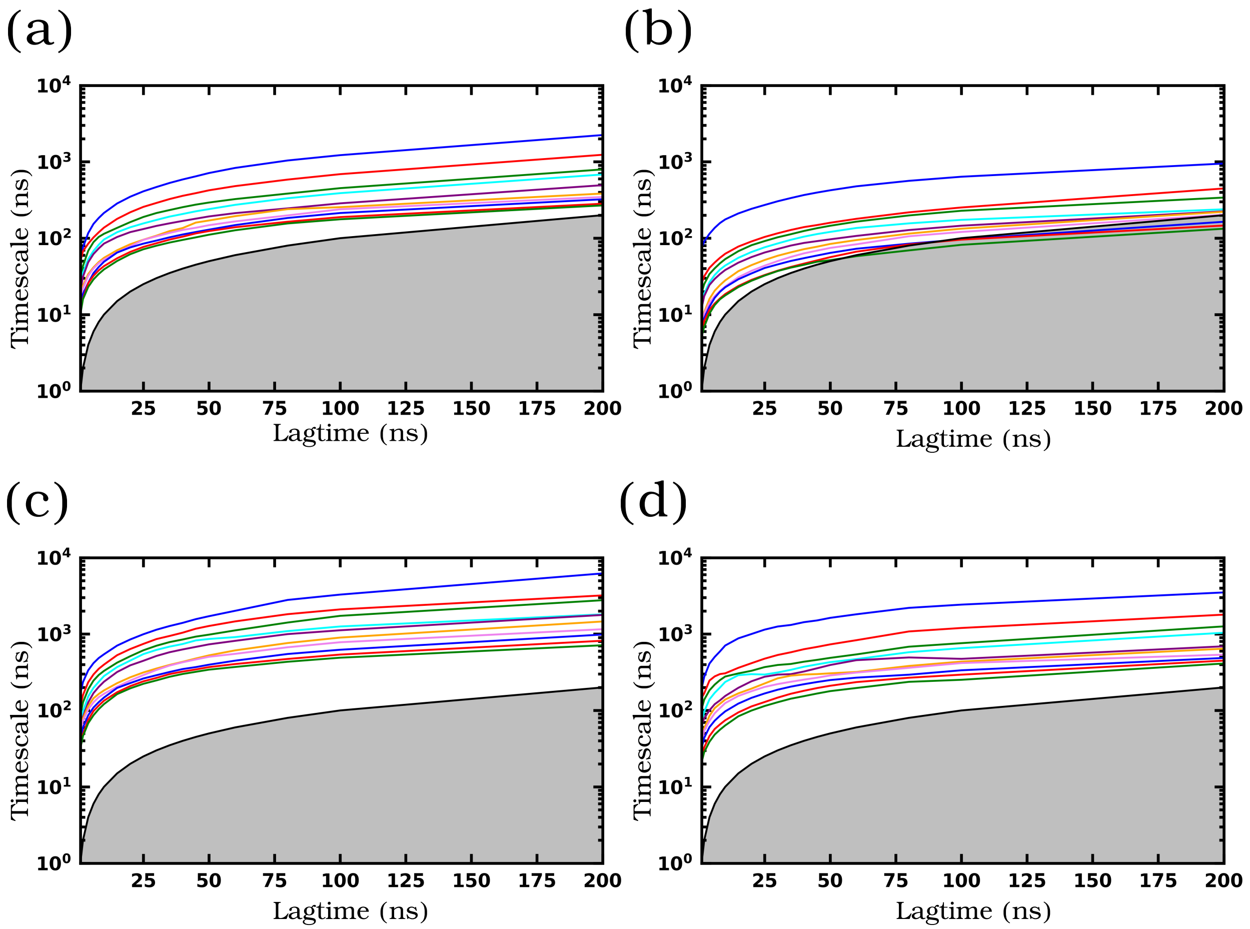


**Figure S9:** ITS of $\alpha$S for (a) Non-annealed cosine VAE (b) Non-annealed logistic VAE (c) Cosine annealed VAE and (d) Logistic annealed VAE.

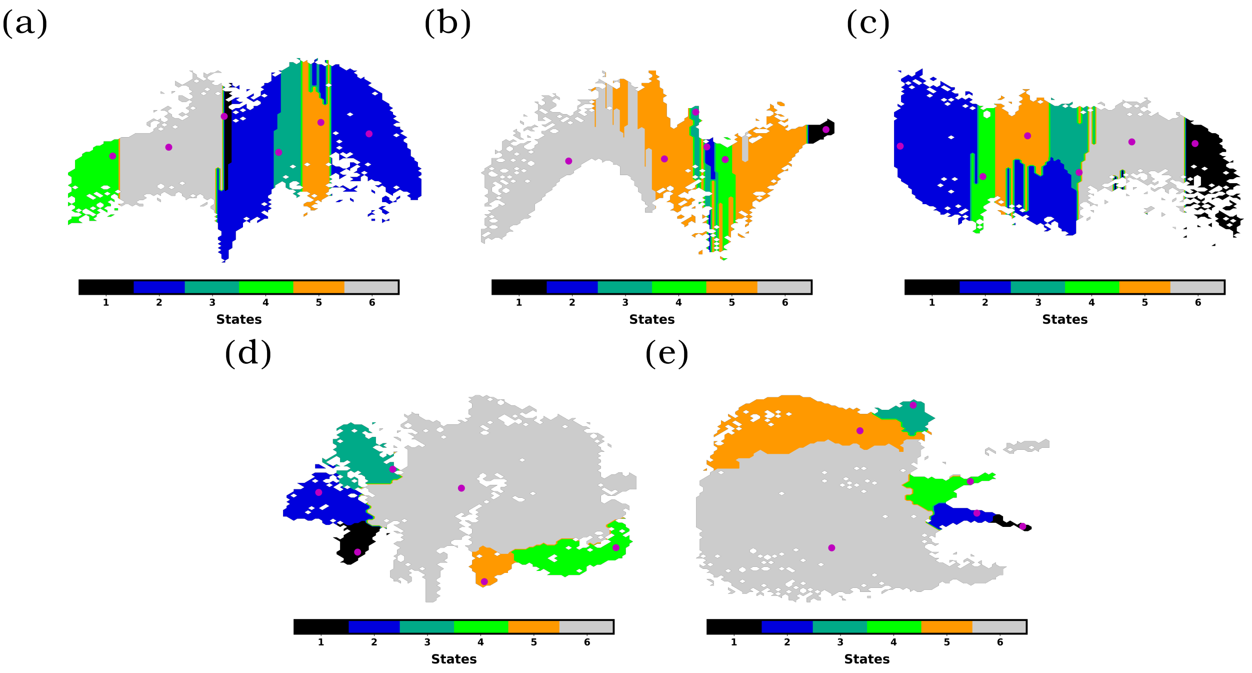


**Figure S10:** MSM macrostates of $\alpha$S for (a) Non-annealed linear VAE (b) Non-annealed cosine VAE (c) Non-annealed logistic VAE (d) Cosine annealed VAE and (e) Logistic annealed VAE.

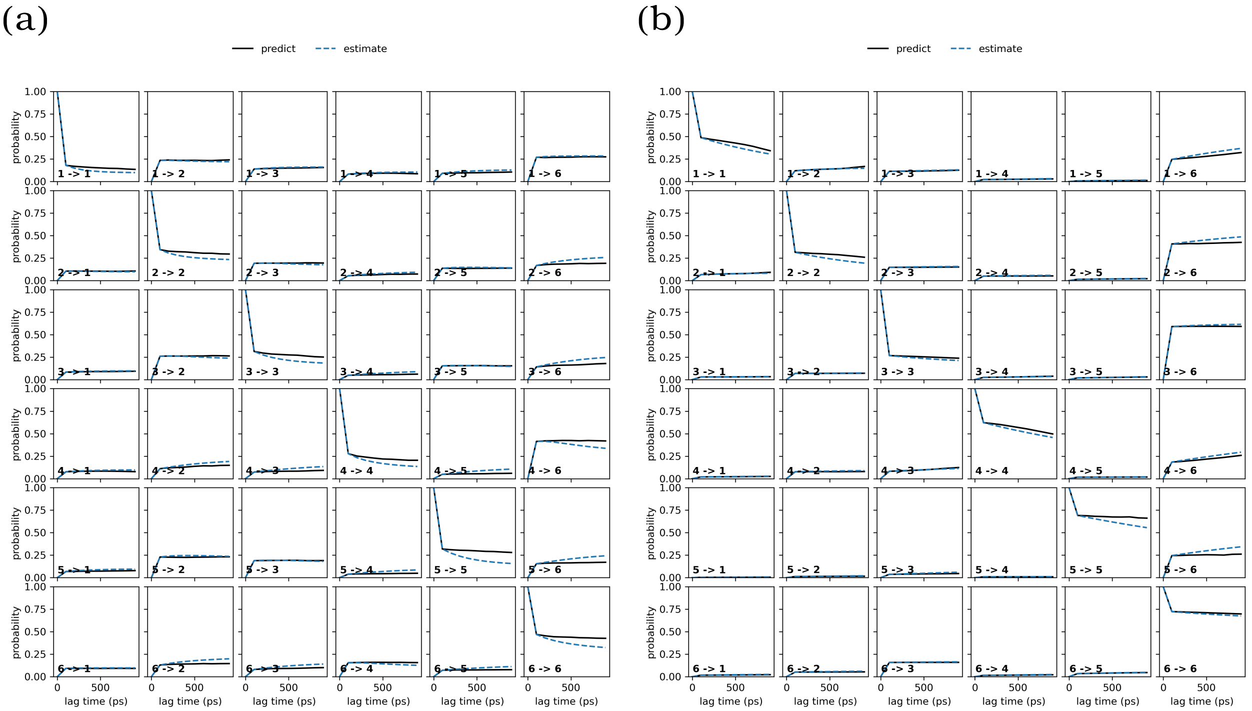


**Figure S11:** CK test of MSM macrostates of $\alpha$S for (a) Non-annealed linear VAE (b) Linear annealed VAE.

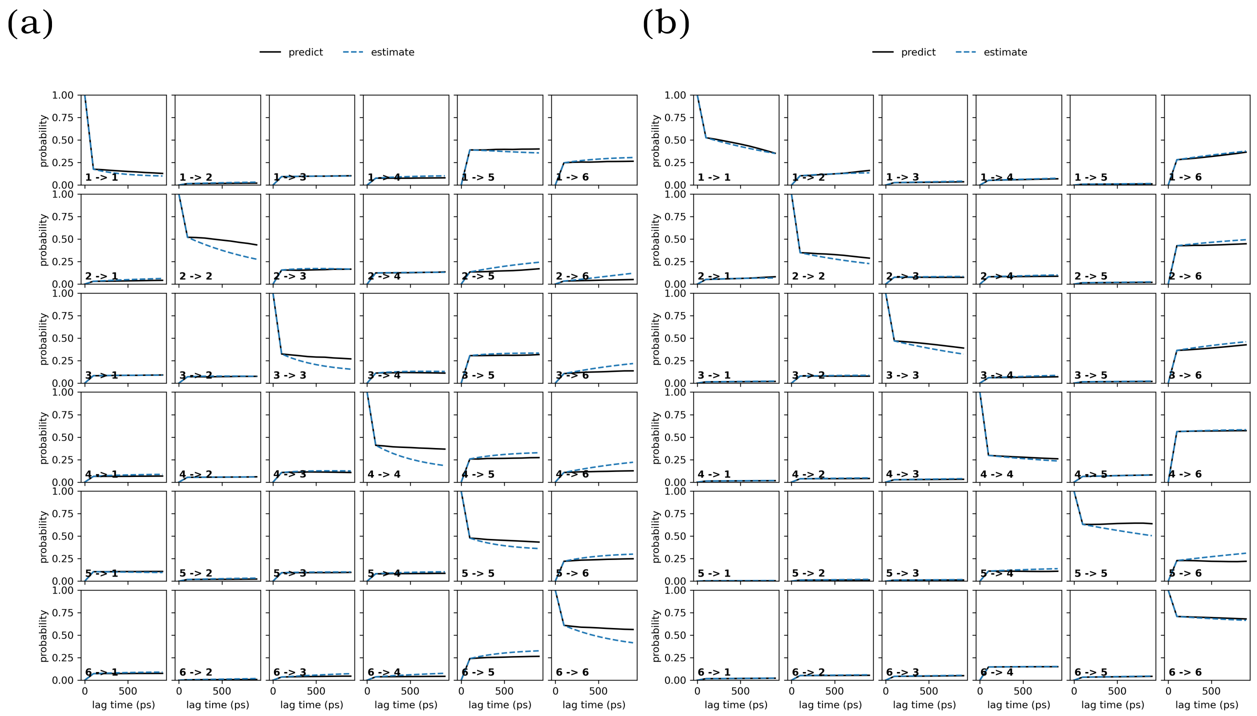


**Figure S12:** CK test of MSM macrostates of $\alpha$S for (a) Non-annealed cosine VAE (b) Cosine annealed VAE.

(a) (b)

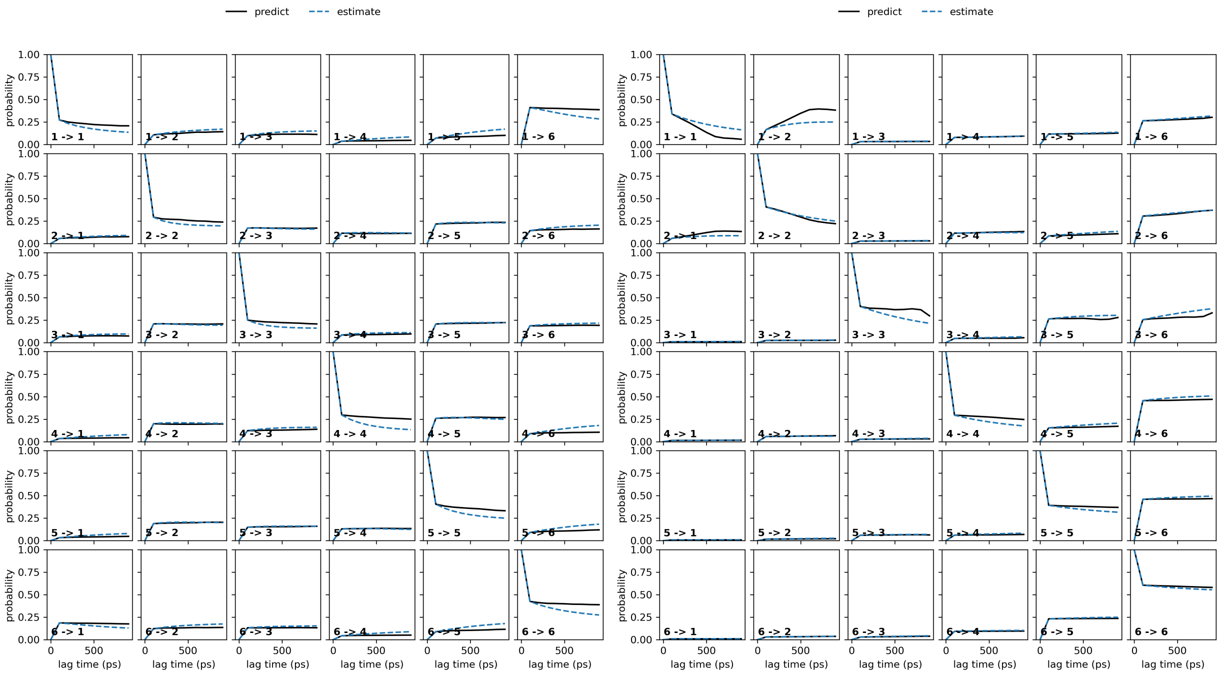


**Figure S13:** CK test of MSM macrostates of $\alpha$S for (a) Non-annealed logistic VAE (b) Logistic annealed VAE.

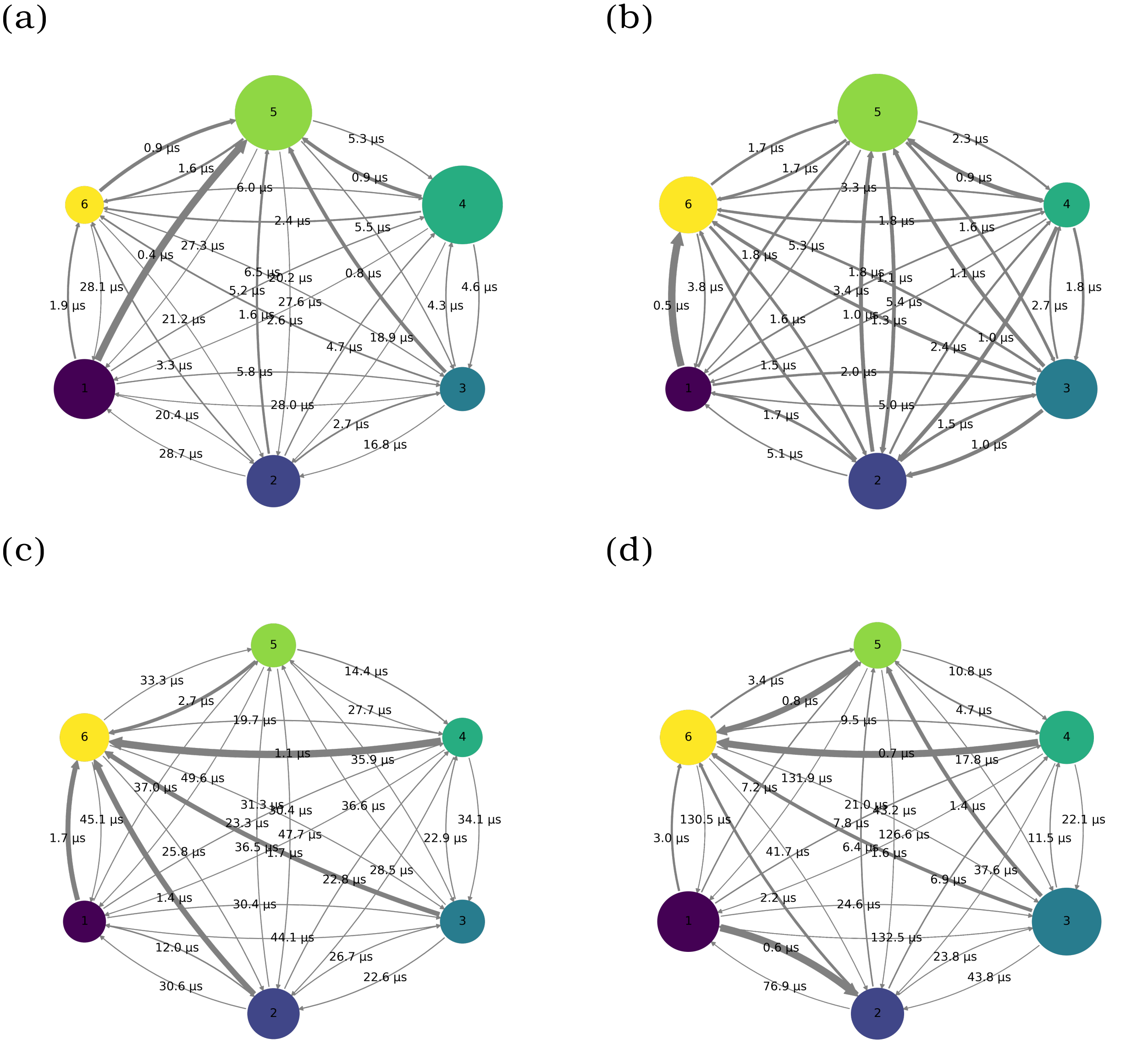


**Figure S14:** MFPT among the MSM macrostates of $\alpha$S for (a) Non-annealed cosine VAE (b) Non-annealed logistic VAE (c) Cosine annealed VAE and (d) Logistic annealed VAE.
